# The complex assembly of N-acetyl-L-glutamate kinase with the PII signal transduction protein follows a two-step mechanism

**DOI:** 10.64898/2026.08.10.743959

**Authors:** Abdalla A. Elshereef, Dmitry Shvarev, Khaled A. Selim

## Abstract

PII signal transduction proteins control central carbon/nitrogen metabolism via interacting with and modulating many cellular targets. In photoautotrophs, N-acetyl-L-glutamate kinase (NAGK), the rate-limiting enzyme of arginine biosynthesis, is the primary target of PII. Here, we focused on how PII initiates the interaction with NAGK. Through biochemical and biophysical analyses, we show that the PII variant lacking the T-loop (PII_ΔT-loop_) is blocked in the first association step with NAGK, but is still able to form a stable complex. However, the NAGK interaction with PII_ΔT-loop_ is weaker than with the wild-type PII as indicated by a decrease in the affinity. Using single particle cryo-EM, we resolved the structure of the encounter PII_ΔT-loop_-NAGK complex, revealing that the PII_ΔT-loop_-NAGK interface is established by the B-loop residues of PII. Thus, our data indicate a two-step process of PII-NAGK complex formation with B-loop initiating the contact with NAGK followed by the insertion of the PII T-loop deeply into the NAGK cavity to fully activate the enzyme.

## INTRODUCTION

PII proteins form an ancient superfamily of signal transduction proteins widespread in all domains of life from prokaryotes to eukaryotes, including bacteria, archaea, and plants.^1–3^ Structurally, PII signal transduction proteins are characterized by a highly conserved trimeric architecture with a ferredoxin-like fold and three distinctive loop regions (T-, B-and C-loops).^2–4^ The loop regions are located near the three intercommunicating metabolite binding sites, located at the clefts between the three subunits, and play a major role in intramolecular signaling, ligand binding, and protein-protein interactions.^1,2^ Through interdependent binding to ATP, ADP, or citric acid cycle-metabolite (2-oxoglutarate; 2-OG), PII proteins function as cellular sensors for energy, carbon, and nitrogen status of the cell.^1,2,5,6^ Based on the ligation state, the surface-exposed T-loop (residues 38-54), which protrudes from each subunit of the PII trimer, contributes to the functional diversity of PII interactions by adopting different conformations.^2^ This allows PII proteins to regulate and bind to various cellular receptors, such as transcription factors, metabolic enzymes, transporters, and small regulatory proteins.^1,2,7–9^ Thus, PII proteins are considered the major regulatory hubs of cellular metabolism.^2,3^

In photoautotrophs, the primary PII target is the N-acetyl-L-glutamate (NAG) kinase (NAGK), the rate-limiting enzyme of arginine (Arg) biosynthesis.^9–12^ NAGK catalyzes the ATP-dependent phosphorylation of NAG to form N-acetylglutamyl-5-phosphate, which is subsequently converted to ornithine and ultimately to arginine within the pathway.^9–12^ The PII-NAGK interaction is highly conserved among cyanobacteria, algae and plants, supporting an essential role for PII signaling in controlling Arg homeostasis.^1,2,10^ Even thought to be restricted to photoautotrophs, a recent finding suggested that the PII-NAGK interaction is also conserved among heterotrophic bacteria.^7,13^ The enzymatic activity of NAGK is feedback inhibited by Arg and enhanced in a complex with PII.^6,9–12^ At sufficient intracellular nitrogen concentrations, indicated by low 2-OG levels, PII forms an ATP-dependent activating complex with NAGK^14^, which additionally diminishes the feedback inhibitory effect of Arg on the NAGK.^6,10–12,15^ Under poor nitrogen supply, indicated by high 2-OG levels^1^, the 2-OG binding to PII causes strong conformational changes in the T-loop^5^, which in turn impairs the PII-NAGK interaction.^6,10–12,15,16^

Structurally, NAGK forms a symmetric hexameric toroid ring composed of three dimers, in which both faces of the toroid are sandwiched by a PII trimer from the top and bottom.^12,14,17^ Each monomer of the PII trimer interacts with the adjacent NAGK monomer^1,10^, whereas the T-loop inserts deeply into the NAGK interdomain cavity. Especially the distal residues of the T-loop (amino acids 42-52) undergo strong rearrangements enabling its deep insertion into NAGK by adapting a compact conformation.^14,17^ The T-loop residues R45, R47, G48, S49, and E50 play key roles in stabilizing the PII-NAGK interaction through stabilizing the PII T-loop compact conformation, needed for the interaction with NAGK.^14^ Consequently, these interactions between PII and NAGK cause a pronounced conformational change in NAGK that alters its enzymatic properties. First, the allosteric Arg-binding sites are enlarged, decreasing the NAGK affinity for Arg and thus elevating the inhibitory Arg concentration. Second, the PII-NAGK complex formation increases the affinity for the substrate NAG, thus enhancing the overall catalytic efficiency of NAGK.^14,17^ Recently, we showed that a tight PII-NAGK complex formation is required to relieve NAGK from Arg feedback inhibition, however, the weak complex is sufficient to activate the NAGK in the absence of Arg.^11^ Nevertheless, despite all those efforts to elucidate the impact of PII on NAGK activity, the molecular details of the PII-NAGK complex association/dissociation are still lacking.

Notably, it was shown that the I86N substitution in the other functional loop of PII, the B-loop, causes the formation of a hydrogen bond with T43 of PII (PDB: 2XBP).^16^ This forces the PII T-loop to adopt a compact, bent conformation similar to the T-loop conformation in the PII-NAGK complex.^14,16^ Consequently, the PII_(I86N)_ variant seems to bind constitutively to NAGK *in vitro* and *in vivo*.^16,18^ Based on this observation, it was proposed that the PII-NAGK complex assembly follows a two-step mechanism^16^: first, an encounter complex involving the B-loop of PII is formed. This includes a contact between R233 of NAGK and E85 of PII, which initiates the bending of the extended T-loop. Also, this interaction breaks a salt bridge between E85 and R47 in the PII T-loop, which is now free and can adopt the bent structure. The bent conformation of the T-loop is stabilized by a new salt bridge between K58 and T-loop residue E44. In the second step, the bent T-loop inserts deeply into the NAGK clefts to form the tight PII-NAGK complex.^14^

To test this hypothesis experimentally, we aimed to capture the first encounter PII-NAGK complex by creating a truncated T-loop version of PII (PII_ΔT-loop_; lacking residues 47-53), which should stay permanently in the first-step contact with NAGK and would be unable to form the stable bent T-loop conformation. Moreover, in response to the metabolic status of the cell, we examined the roles of various effector molecules (NAG, ATP, ADP, 2-OG and Arg) on the dynamics of PII_ΔT-loop_-NAGK interaction and the overall catalytic activity. Finally, we isolated the encounter PII_ΔT-loop_-NAGK complex and solved its structure using single particle cryo-EM to unravel the molecular details of PII_ΔT-loop_-NAGK complex assembly and PII-induced NAGK activation.

## RESULTS

### PII_ΔT-loop_ variant forms the encounter PII-NAGK complex independent of effector molecules

The formation of PII-NAGK complex requires a compact bending of the protruding PII T-loop and its insertion into the interdomain cleft of the NAGK subunits for anchoring PII protein.^14^ Before the deep insertion of the PII T-loop into the NAGK cavity, it was proposed that an initial encounter PII-NAGK complex forms. This complex involves direct contact between E85 of the PII B-loop and R233 of NAGK, allowing the T-loop to adopt a bent conformation that interacts with NAGK via its residues 44-50.^16^ Thus, the distal T-loop residues (44-50) play dual roles by forming a strong network of interactions with NAGK and additionally stabilizing the PII T-loop in the compact bended conformation.^14,16,17^ To examine this hypothesis of the two-step mechanism of PII-NAGK complex formation, we created a PII variant lacking the essential T-loop residues 47-53 (PII_ΔT-loop_).^19^ In the PII_ΔT-loop_ variant, the bent T-loop conformation cannot form, and therefore the PII-NAGK complex formation is expected to be blocked in the first step. For this purpose, we heterologously expressed and purified the recombinant wild-type PII (PII_WT_) and its PII_ΔT-loop_ variant as C-terminal His_8_-tagged^19^, while NAGK was produced as Strep-tagged.

To test the direct interaction between NAGK and PII variants (PII_WT_ and PII_ΔT-loop_), and to assess the influence of known PII or NAGK effector molecules on the complex assembly/dissociation, we used biolayer interferometry (BLI).^20^ In the BLI experiments, the His-tagged PII proteins (either PII_WT_ or PII_ΔT-loop_) were immobilized on a Ni^2+^-sensors. Strep-tagged NAGK was then sequentially injected at varying concentrations (31-1000 nM), either in the presence or absence of saturating concentration of PII effectors (ATP, ADP, or 2-OG; all at 1 mM) or NAGK effectors (50 mM NAG and 100 µM Arg) to monitor the difference in the response units (λ in nm) and calculate the binding kinetics of PII-NAGK complex formation^6,9,16,21^ (Table 1).

**Table 1.**
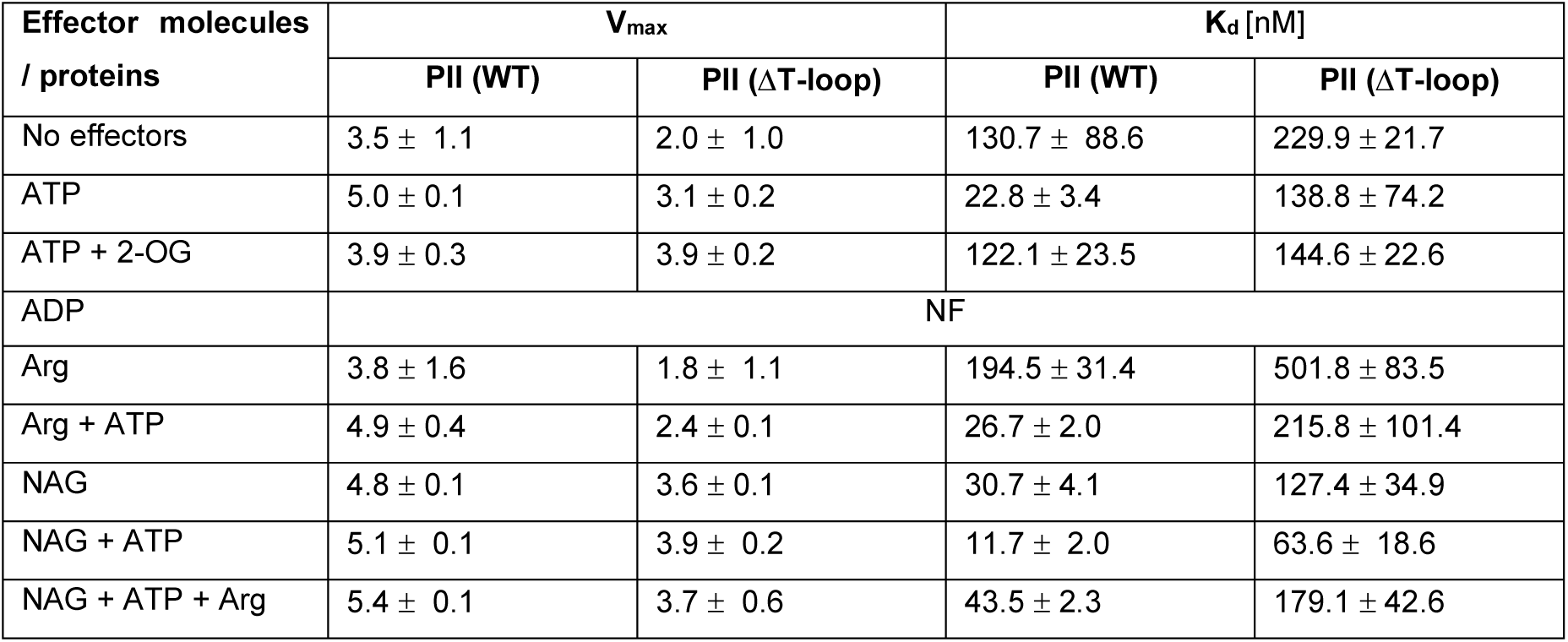
Kinetics parameters of PII_WT_-NAGK or PII_ΔT-loop_-NAGK interactions as revealed by BLI analysis. NF: not fitted.

| Effector molecules<br>/ proteins | V<sub>max</sub> |  | K<sub>d</sub> [nM] |  |
| --- | --- | --- | --- | --- |
|  | PII (WT) | PII (ΔT-loop) | PII (WT) | PII (ΔT-loop) |
| No effectors | 3.5 ± 1.1 | 2.0 ± 1.0 | 130.7 ± 88.6 | 229.9 ± 21.7 |
| ATP | 5.0 ± 0.1 | 3.1 ± 0.2 | 22.8 ± 3.4 | 138.8 ± 74.2 |
| ATP + 2-OG | 3.9 ± 0.3 | 3.9 ± 0.2 | 122.1 ± 23.5 | 144.6 ± 22.6 |
| ADP | NF |  |  |  |
| Arg | 3.8 ± 1.6 | 1.8 ± 1.1 | 194.5 ± 31.4 | 501.8 ± 83.5 |
| Arg + ATP | 4.9 ± 0.4 | 2.4 ± 0.1 | 26.7 ± 2.0 | 215.8 ± 101.4 |
| NAG | 4.8 ± 0.1 | 3.6 ± 0.1 | 30.7 ± 4.1 | 127.4 ± 34.9 |
| NAG + ATP | 5.1 ± 0.1 | 3.9 ± 0.2 | 11.7 ± 2.0 | 63.6 ± 18.6 |
| NAG + ATP + Arg | 5.4 ± 0.1 | 3.7 ± 0.6 | 43.5 ± 2.3 | 179.1 ± 42.6 |

In the absence of any effector molecules, the PII_ΔT-loop_ variant was still able to bind NAGK, albeit with significantly lower affinity, as indicated by higher K_d_ value and lower V_max_ of λ response compared to PII_WT_ (Figure 1A). This represents a nearly two-fold decrease in binding affinity, as evidenced by a doubling of the K_d_ for the PII_ΔT-loop_-NAGK complex assembly compared to PII_WT_ protein (Table 1). In the presence of ATP, which is known to stabilize and promote the PII-NAGK interaction^12,15^, PII_WT_ formed a very stable complex with NAGK as indicated by a sharp decrease in the K_d_ value compared to the apo complex. Also, the PII_WT_-NAGK complex did not dissociate from the BLI-sensors after the end of the injection phase (compare Figure 1B with 1A). For the PII_ΔT-loop_ variant, the ATP only slightly enhanced the binding kinetics of PII_ΔT-loop_ to NAGK (compare Figure 1B with 1A), indicating a weaker complex formation compared to PII_WT_. Additionally, the PII_ΔT-loop_-NAGK complex was not stable and dissociated quickly compared to PII_WT_ (Figure 1B), implying that this variant additionally does not sense ATP properly.

**Figure 1.**
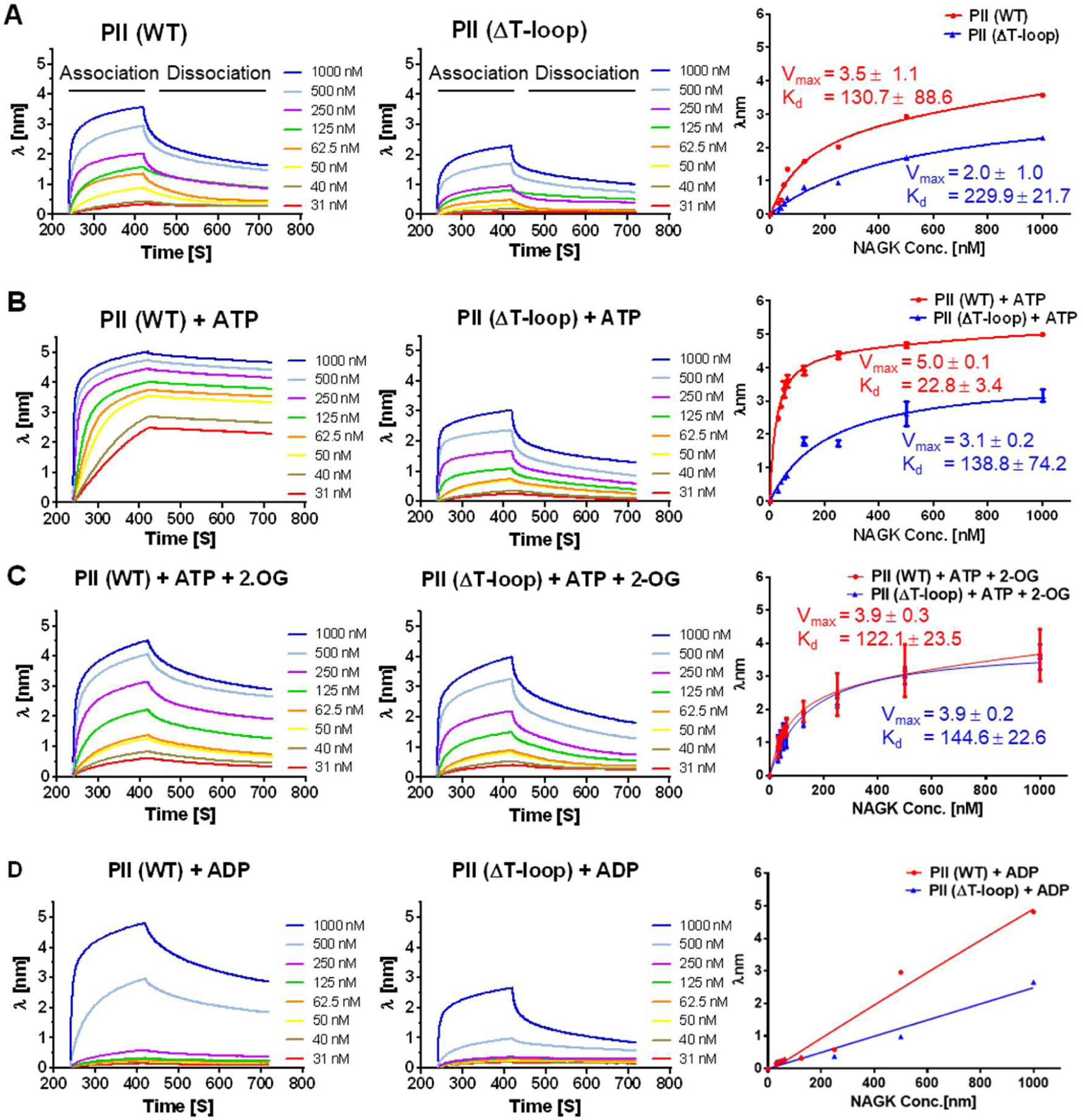
BLI analysis of the impact of PII effector molecules on PII_WT_-NAGK or PII_ΔT-loop_-NAGK interactions. Assays were performed in the absence (A) or presence of PII effector molecules: ATP (B), 2-OG (C), or ADP (D). Graphs display the association and dissociation phases of PII_WT_-NAGK or PII_ΔT-loop_-NAGK complexes at varying NAGK concentrations, as indicated. The left and middle panels show the interaction of PII_WT_-NAGK and PII_ΔT-loop_-NAGK, respectively. The right panels show the V_max_ and K_d_ values for the assembly of PII_WT_-NAGK and PII_ΔT-loop_-NAGK complexes, calculated using a one-site binding model (except in the presence of ADP, which could not be fitted). The V_max_ and K_d_ values are given as the mean ± standard deviation of at least 2-3 independent experiments.

Furthermore, we tested the impact of ADP and 2-OG, which are known to have a negative effect on PII-NAGK interaction by promoting complex dissociation.^5,15,21^ Indeed, in contrast to ATP, in the presence of 2-OG, the stability of the PII_WT_-NAGK complex was weakened, as indicated by an increase in the K_d_ value and immediate dissociation after the end of the injection phase (compare Figure 1C with 1B). On the other hand, 2-OG did not affect the PII_ΔT-loop_-NAGK complex as demonstrated by kinetic parameters comparable to those of the PII_WT_-NAGK complex (Figure 1C) and the PII_ΔT-loop_-NAGK complex in the presence of ATP alone (compare K_d_ values in Table 1, and Figure 1C with 1B). These results indicate that the PII_ΔT-loop_ variant lost its ability to sense 2-OG, consistent with the T-loop’s role in stabilizing the 2-OG interaction. Next, we analyzed the effect of ADP on the complex assembly. In the presence of ADP, the complex was unstable and assembled only under high NAGK concentrations and rapidly dissociated (Figure 1D), which hampered the calculation of kinetic constants. NAGK showed an even weaker association to the PII_ΔT-loop_ variant than with PII_WT_, as indicated by a lower λ response (Figure 1D). Altogether, these results further support that the PII_ΔT-loop_ protein forms a weaker complex with NAGK compared to the PII_WT_ protein, and the PII_ΔT-loop_-NAGK complex is likely blocked at the first step of the complex assembly due to the absence of a functional T-loop; therefore, it dissociates faster (e.g., Figure 1B).

Next, we wanted to investigate the influence of NAGK effector molecules (its substrate NAG and its inhibitor Arg) on the complex assembly, which had not been previously tested. First, we examined the effect of Arg on the PII-NAGK complex formation. In the presence of 100 µM Arg, both assembly and dissociation of PII-NAGK complex were strongly impaired (compare Figure 2A with Figure 1). This was indicated by higher K_d_ values (Table 1) compared to, for example, complex formation in the presence of ATP or the absence of any effector molecules (compare Figure 2A with Figure 1A,B). Again, the affinity and V_max_ values of the PII_ΔT-loop_ variant were nearly two-fold lower than those of the PII_WT_ protein (Figure 2A and Table 1). This result implies that Arg acts as an inhibitor of the complex assembly in a similar manner to 2-OG and ADP (Figure 1C,D). The addition of ATP to the binding assays relieved the inhibitory effect of Arg on the complex by preventing its fast dissociation, and hence enhanced the affinity of NAGK for both the PII_WT_ and PII_ΔT-loop_ proteins (Table 1 and Figure 2B), and slightly enhanced the V_max_ of the λ response (compare Figure 2B with Figure 2A). Nevertheless, in the presence of Arg and ATP, the PII_WT_-NAGK complex was less stable and dissociated at a faster rate compared to when ATP was present alone (compare the dissociation phases of Figure 2B with Figure 1B), but the association showed comparable affinities as indicated by similar K_d_ and V_max_ values (Table 1). In contrast, the effect of Arg on PII_ΔT-loop_ variant even in presence of ATP was more pronounced, as indicated by a lower V_max_ and an increased K_d_ value compared to ATP alone (compare Figure 2B with Figure 1B and Table 1).

**Figure 2.**
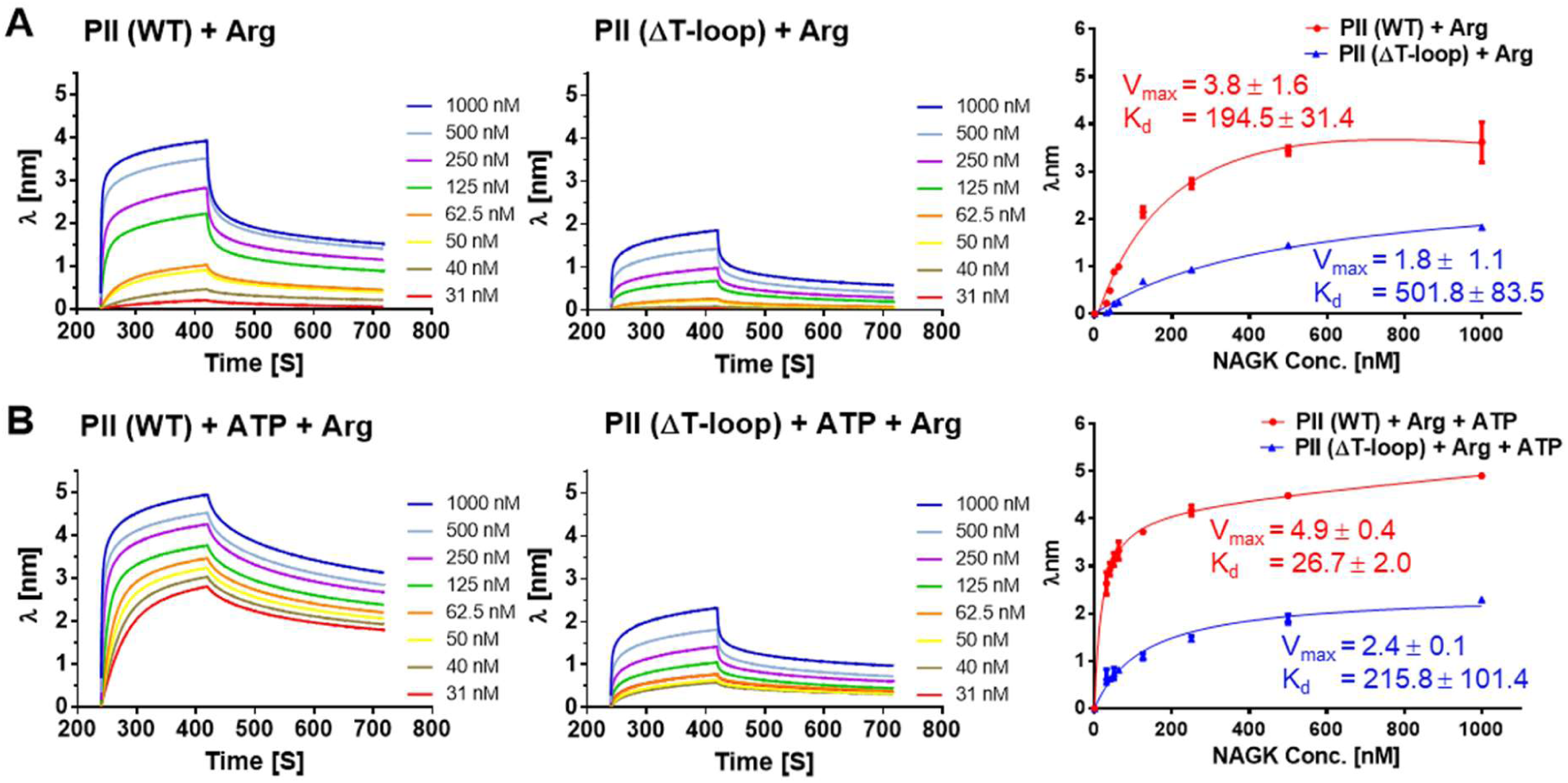
Arg impact on PII_WT_-NAGK or PII_ΔT-loop_-NAGK interactions as revealed by BLI analysis. Assays were performed in the presence of Arg only (A) or in a combination of Arg with ATP (B). Graphs display the association and dissociation phases of PII_WT_-NAGK or PII_ΔT-loop_-NAGK complexes at varying concentrations of NAGK, as indicated. The left and middle panels show the interaction of PII_WT_-NAGK and PII_ΔT-loop_-NAGK, respectively. The right panels show the V_max_ and K_d_ values for PII_WT_-NAGK and PII_ΔT-loop_-NAGK complexes assembly, calculate using a one-site binding model. The V_max_ and K_d_ values are given as the mean ± standard deviation of at least 2-3 independent experiments.

Finally, we tested the effect of NAG on the PII-NAGK complex formation. Surprisingly, NAG prevented the fast dissociation of the complex after the end of the injection phase (compare Figure 3A with Figure 1A). It also enhanced the binding affinity of NAGK toward both PII_WT_ and PII_ΔT-loop_ proteins, as evidenced by lowering K_d_ values (Table 1 and compare Figure 1A with Figure 3A), which were comparable to those observed in the presence of ATP alone (Table 1 and compare Figure 3A with Figure 1B). This indicates that NAG acts like ATP in promoting PII-NAGK complex formation. In the presence of both ATP and NAG, binding affinities were enhanced for both PII variants, and the ATP stabilized the complex further by preventing dissociation, particularly for the PII_WT_ protein (compare Figure 3B with Figures 1B and 3A). Although the PII_ΔT-loop_ still exhibited weaker binding than the PII_WT_ protein, it achieved its strongest affinity under this combined condition, as indicated by its lowest K_d_ value (Table 1 and Figure 3B). In a binding assay that combined all effectors (ATP, NAG and Arg), thereby turning NAGK in a catalytically active state that accumulates its product NAG-phosphate (NAG-P), the PII_ΔT-loop_-NAGK complex showed weaker K_d_ compared to the wild-type complex (Figure 3C). Nevertheless, both PII_WT_-NAGK and PII_ΔT-loop_-NAGK complexes were significantly more stable than in the presence of Arg alone (Figure 2A), further confirming the positive impact of ATP and NAG in enhancing the complex stability and relieving the inhibitory effect of Arg.

**Figure 3.**
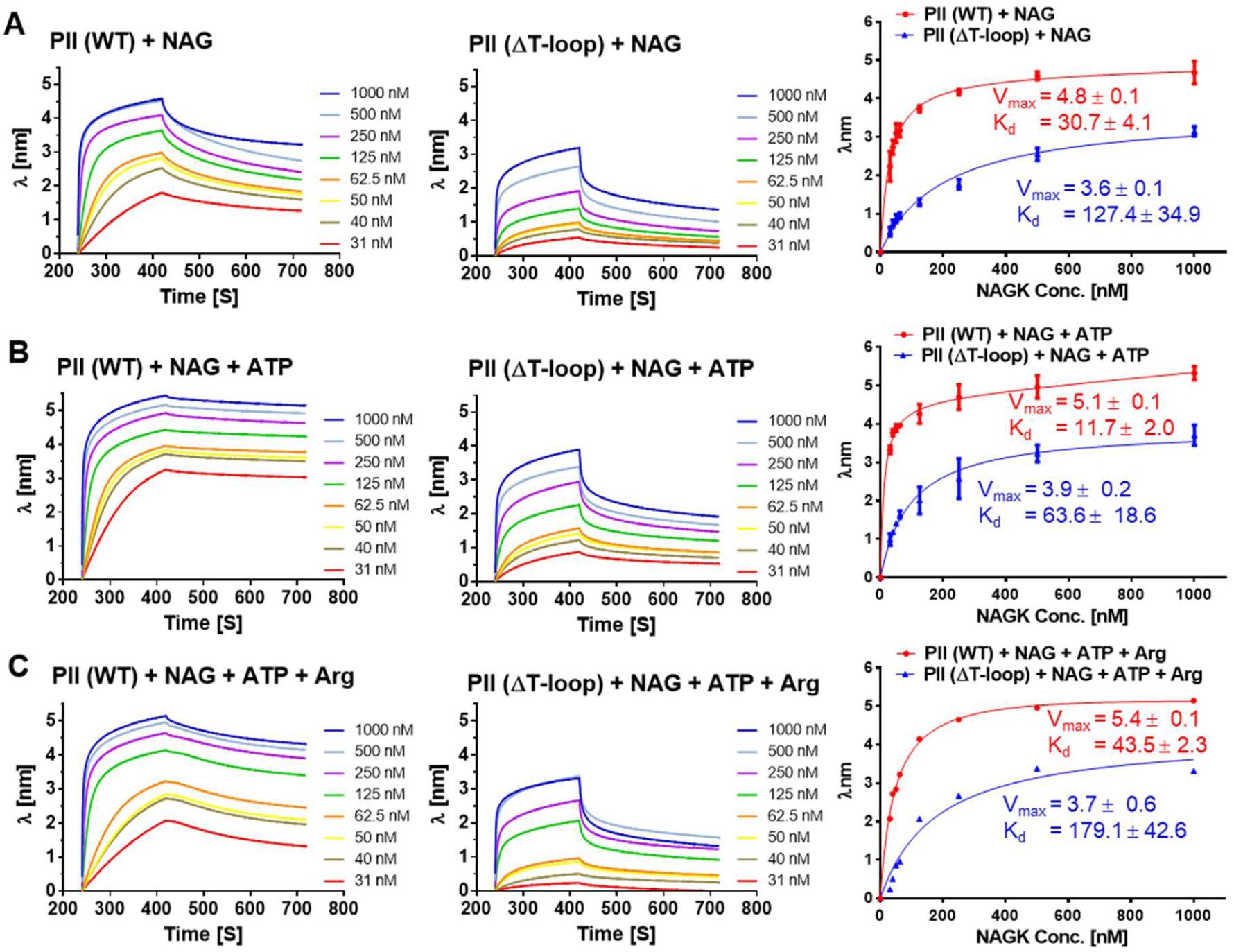
NAG impact on PII_WT_-NAGK or PII_ΔT-loop_-NAGK interactions as revealed by BLI analysis. The assays were performed in the presence of NAG only (A) or in a combination of NAG with ATP (B), or Arg + ATP (C). Graphs display the association and dissociation phases of PII_WT_-NAGK or PII_ΔT-loop_-NAGK complexes at varying concentrations of NAGK, as indicated. The left and middle panels show the interactions of PII_WT_-NAGK and PII_ΔT-loop_-NAGK, respectively. The right panels show the V_max_ and K_d_ values for PII_WT_-NAGK and PII_ΔT-loop_-NAGK complexes assembly, calculated using a one-site binding model. The V_max_ and K_d_ values are given as the mean ± standard deviation of at least 2-3 independent experiments.

Collectively, our protein-protein interaction experiments revealed that the PII_ΔT-loop_ variant has a weaker affinity to NAGK, with an approximately two-fold reduction in kinetic parameters compared to PII_WT_ across all tested conditions (Figures 1-3). This is consistent with the proposed two-step mechanism for the complex assembly, suggesting that the PII_ΔT-loop_ variant is likely trapped in the less stable first-encounter complex and dissociates faster than the wild-type complex.

### PII_ΔT-loop_ variant activates NAGK, however a tight complex is needed to alleviate Arg feedback inhibition

Next, we used a NAGK enzymatic assay to assess whether the weak ability of PII_ΔT-loop_ variant to form a complex with NAGK correlates with its overall catalytic activity compared to that of the PII_WT_ protein.^9,11,12,15^ The kinetic constants for NAGK were determined using NAG as a variable substrate in the presence or absence of PII proteins (PII_WT_ or PII_ΔT-loop_). The PII_ΔT-loop_ variant activated NAGK (Figure 4A), but to a lesser extent than PII_WT_, as indicated by a lower V_max_ and an approximately two-fold increase of the K_m_ for the PII_ΔT-loop_ variant (3.5 ± 0.5 mM). The changes in NAGK kinetic parameters triggered by PII_ΔT-loop_ variant indicate that it is still able to interact with NAGK, but seemingly transiently, compared to the PII_WT_ protein.

**Figure 4.**
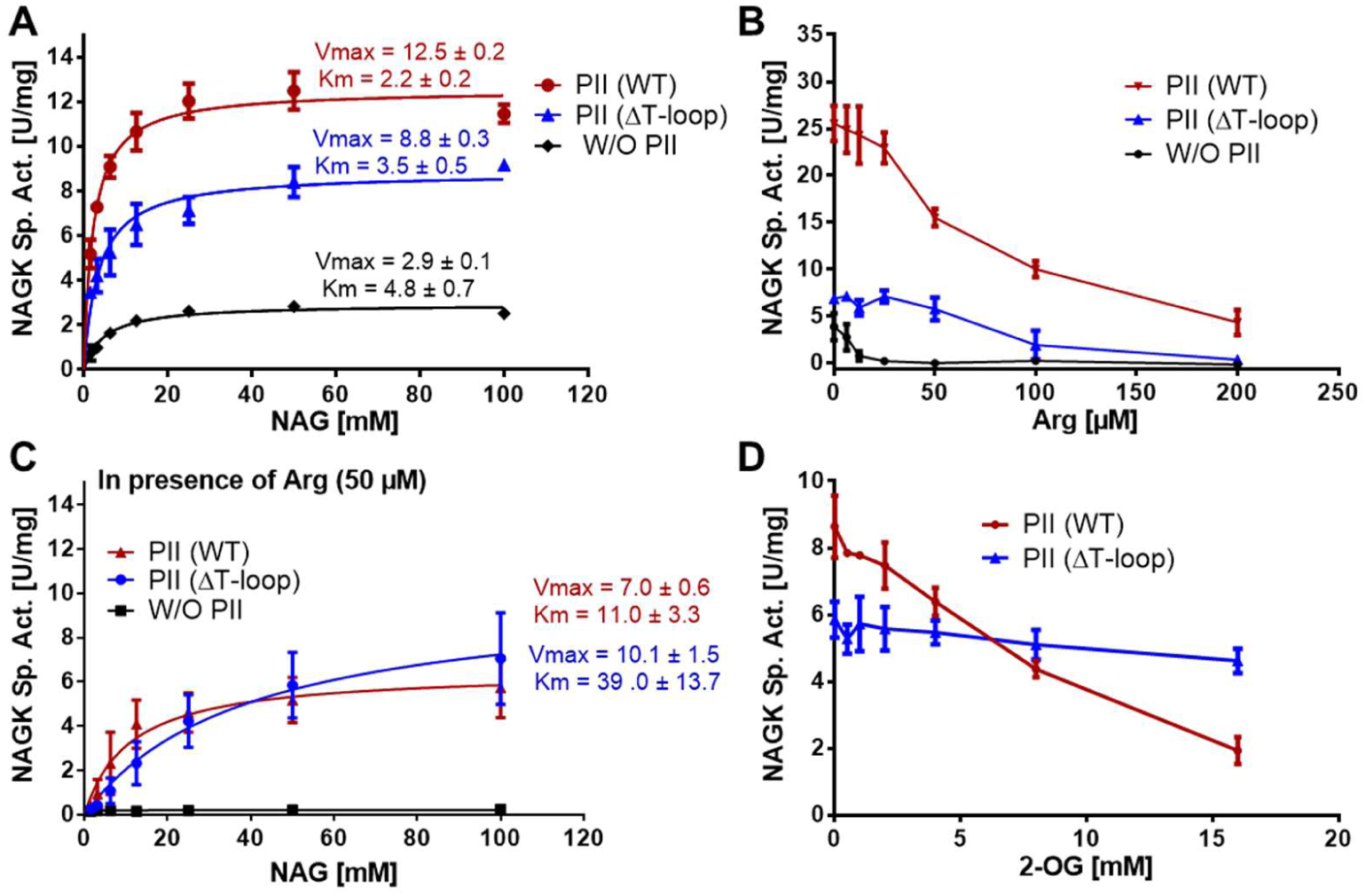
Impact of PII_WT_ and PII_ΔT-loop_ variants on NAGK enzymatic activity. Graphs depict NAGK activity with and without PII protein variants under different conditions: (A) in presence of NAG as variable substrate concentrations; (B) in presence of Arg as variable inhibitor concentrations and 50 mM fixed concentration of NAG; (C) in presence of NAG as variable substrate concentrations and 50 µM fixed concentration of Arg; (D) in presence of 2-OG as variable inhibitor concentrations and fixed concentration of NAG (50 mM) and Arg (50 µM). Kinetic parameters were calculated using GraphPad Prism software. The error bars represent the standard deviation (SD) of biological triplicates (n=3).

The alleviation of NAGK from Arg feedback inhibition through PII-NAGK complex formation is the rate-limiting step for the metabolic switch of the Arg biosynthetic pathway.^6,11,12,15^ To examine whether the PII_ΔT-loop_ variant relieves NAGK from Arg inhibition, we assessed NAGK activity at a fixed concentration of NAG (50 mM) in the presence of varying concentrations of Arg, both with and without PII variants (PII_WT_ and PII_ΔT-loop_; Figure 4B). Without PII, the Arg feedback inhibition on NAGK occurred with a half-minimal inhibitory concentration (IC_50_) of 8.2 µM (Figure 4B). As expected, the Arg inhibitory effect on NAGK was mitigated in the presence of PII_WT_, yielding an estimated IC_50_ of 125.8 µM (Figure 4B). The PII_ΔT-loop_ variant was still able to partially activate NAGK and relieve it from Arg feedback inhibition. However, the transient PII_ΔT-loop_-NAGK complex was more sensitive to Arg, with an IC_50_ (75.6 µM) approximately half of that of the PII_WT_-NAGK complex (Figure 4B). This further highlights the importance of the PII T-loop-dependent formation of a stable complex with NAGK to alleviate Arg feedback inhibition.

Next, we determined the kinetics of NAGK using NAG as a variable substrate in the presence of 50 µM Arg, comparing with and without PII proteins (PII_WT_ or PII_ΔT-loop_). In the presence of 50 µM Arg, the overall kinetic parameters (Figure 4C) were impaired compared to those in the absence of Arg (Figure 4A). Without PII, NAGK was completely inhibited, while the PII_ΔT-loop_ variant was able to activate NAGK, but to a lesser extent than the WT_WT_ protein, as demonstrated by a higher K_m_ (Figure 4C). These results further support the notion of two-step mechanism for PII-NAGK complex assembly, in which the PII_ΔT-loop_-NAGK complex remains locked in the initial encounter state and is unable to relieve the Arg feedback inhibition as efficiently as PII_WT_.

Finally, we determined the influence of 2-OG on the complex stability and overall NAGK activity, as 2-OG is known as an antagonizing metabolite that disrupts the PII-NAGK interaction. In the presence of 50 µM Arg and 50 mM NAG, 2-OG had a pronounced negative effect on PII_WT_-NAGK activity (with an IC_50_ of 7.2 mM; Figure 4D), antagonizing the protective effect of PII_WT_ on NAGK.^1,10^ However, in the case of the complex formed with the PII_ΔT-loop_ variant, the addition of 2-OG had a negligible effect on the NAGK activity, even at higher 2-OG concentrations (Figure 4D). This further confirms that the PII_ΔT-loop_ variant is unable to bind 2-OG effectively, consistent with our BLI analysis (Figure 1C) and in agreement with the role of the T-loop in anchoring 2-OG.^5^

Our binding and enzymatic assays suggested that the PII_ΔT-loop_ variant is still able to interact with NAGK, presumably due to the formation of the encounter primary complex between PII and NAGK. To further confirm this hypothesis, we isolated PII_ΔT-loop_-NAGK complex using analytical size-exclusion chromatography (SEC) coupled with multi-angle light scattering (MALS) (Figure S1).^9,11^ The PII_WT_ protein is known to form a stable complex with NAGK with a molecular weight of 276 kDa, corresponding to two PII trimers (each trimer of approximately 40.8 kDa) sandwiching one hexameric NAGK (approximately 194 kDa).^11^ The SEC-MALS experiment revealed that both the PII_WT_ and PII_ΔT-loop_ proteins co-eluted with NAGK from the SEC column, with an apparent molecular weight of 265.5 kDa (Figure S1A), which agrees with the theoretical mass of the PII-NAGK complex. NAGK alone appeared at 199.8 kDa (Figure S1A). Of note, the peak of the PII_ΔT-loop_-NAGK complex was approximately half the size of the corresponding PII_WT_-NAGK complex peak. Additionally, the peak of uncomplexed PII_ΔT-loop_ was almost double the size of the uncomplexed PII_WT_ peak. These results suggest that the PII_ΔT-loop_ variant has a tendency to form unstable complex with NAGK that dissociates during SEC (Figure S1A). This experiment further confirmed our BLI analysis (Figures 1-3) and supported the conclusion that the physical interaction between NAGK and the PII_ΔT-loop_ variant is weak due to the formation of the primary encounter PII-NAGK complex.

### In complex with PII_ΔT-loop_, NAGK is bound to Arg and adopts an open conformation

To reveal the molecular basis of the decreased affinity of PII_ΔT-loop_ to NAGK and to explain the lower activity of NAGK in complex with PII_ΔT-loop_, we isolated the encounter PII_ΔT-loop_-NAGK complex (Figure S1). Using cryo-EM, we resolved its D3-symmetric structure to an overall resolution of 3.2 Å in the presence of ATP, NAG, and Arg (Figure 5, Figure S2, and Figure S3). The PII_ΔT-loop_-NAGK complex (Figure 5A) displayed the standard architecture similar to previously published structures of the PII_WT_-NAGK complex^12,14,17^ (Figure S4). The NAGK ring, formed by a trimer of NAGK dimers, is sandwiched between two PII_ΔT-loop_ trimers from top and bottom (Figure 5A). The cryo-EM density allowed us to assign molecules of the added ligands to the PII_ΔT-loop_-NAGK structure (Figure 5B-G and Figure S3D). Each NAGK subunit contains one molecule of Mg^2+^-ATP and NAG close to the enzyme’s active site (Figure 5D-F and Figure S5B), similarly to the homologous NAGK structures.^22^ The main NAGK residues involved in the contacts with Mg^2+^-ATP are K37, D211, D191 and K250, located in α-helices αG and αE and in β-strands β1 and β11 of NAGK (Figure 5F). Moreover, we observed incomplete density corresponding to NAG (Figure S3D), suggesting a partial occupancy of NAGK at this position. NAG is coordinated by NAGK residues R94, N187, N189 (Figure 5F). The observed stabilization of the PII_ΔT-loop_-NAGK complex in such a pre-hydrolysis configuration, where ATP is intact and NAG is not phosphorated, might explain the weak activity of the complex in concordance with the results of the enzymatic assays (Figure 4A,C). Apart from NAG and Mg^2+^-ATP, each NAGK monomer contains a molecule of Arg bound in its expected binding site, close to the N-terminus of NAGK (Figure 5D,G and Figure S5D). The bound Arg is coordinated by residues from α-helices αH and αN and β-strands β11, β15 and β16. These include Y25 and F29, as well as K205, H266, E279 and M289, resembling the Arg-binding site organization observed in homologous proteins.^14,22^

**Figure 5.**
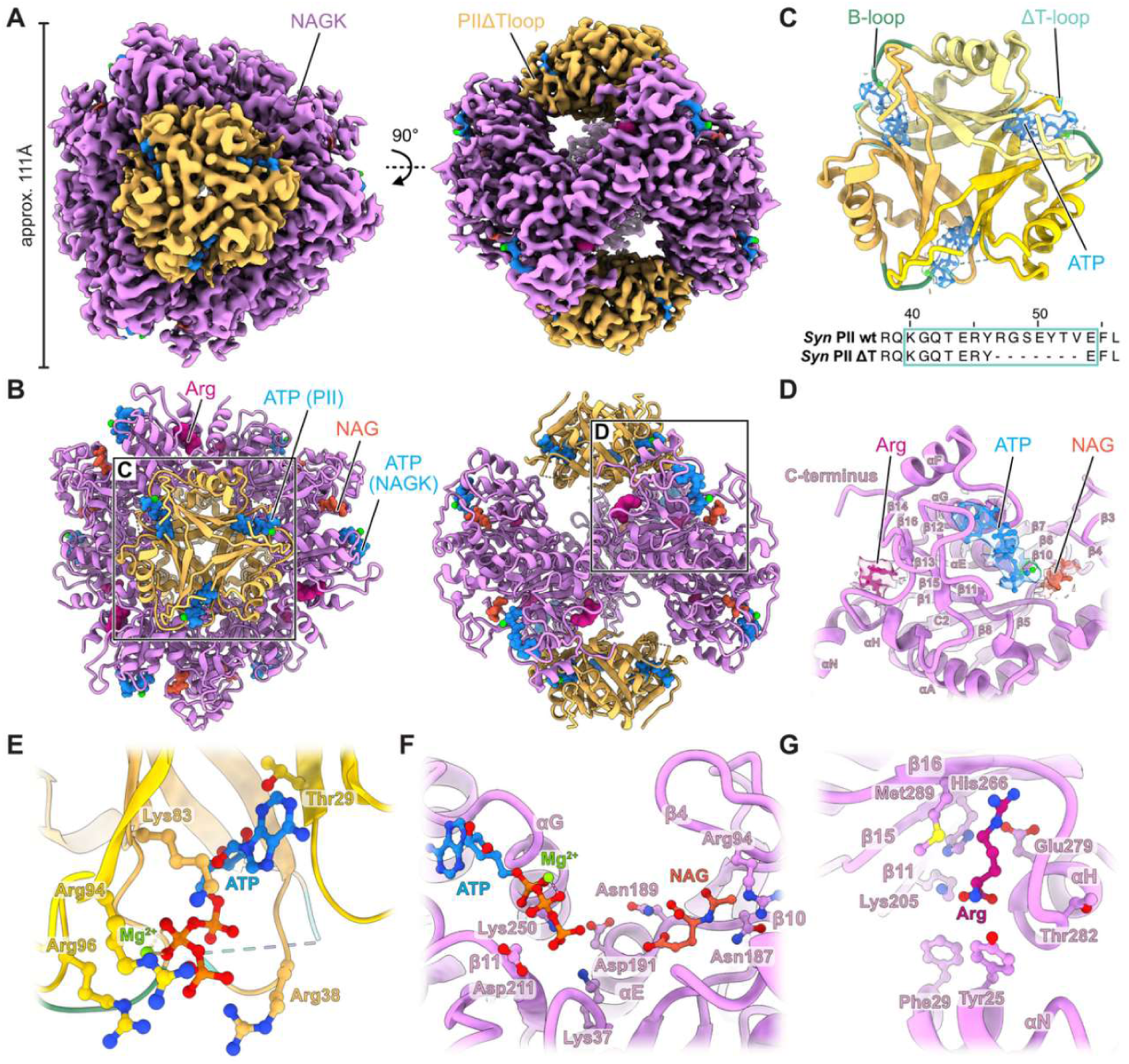
Structure of the PII_ΔT-loop_-NAGK encounter complex. (A) Cryo-EM map of the complex viewed from the PII_ΔT-loop_ side (left) and rotated 90° (right). PII_ΔT-loop_ trimers are colored yellow and NAGK hexamer is purple. (B) Molecular model of the complex fitted into the map. Bound ligands are shown as spheres (Arg in pink; NAG in orange; ATP in dark blue). (C) Ribbon representation of the PII_ΔT-loop_ structure. Individual subunits are colored in different shades of yellow. The characteristic B-loop (green) and truncated T-loop (ΔT-loop in cyan) are shown. Models of bound Mg^2+^-ATP molecules (blue) are represented as balls and sticks, the associated cryo-EM densities are zoned around the models. The sequence alignment of the PII_WT_ and PII_ΔT-loop_ fragments demonstrates the absence of the T-loop in the mutant (the T-loop sequence is marked by the teal box). (D) Overview of NAGK monomer in ribbon representation with bound ligands shown as balls and sticks (coloring as in B). The associated cryo-EM densities are zoned around the ligand models. NAGK α-helices and β-strands are indicated. (E) Close-up view of the PII ATP binding site. ATP (dark blue), Mg^2+^ (lime green) and the surrounding amino acid residues of PII (yellow), involved in the ligand coordination, are indicated. (F) Close-up view of the NAGK ATP and NAG binding site. ATP (dark blue), Mg^2+^ (lime green), NAG (orange) and the surrounding amino acid residues, as well as the α-helices and β-strands of NAGK (purple), involved in the ligand coordination, are indicated. (G) Close-up view of the Arg binding site. Arg (pink) and the surrounding amino acid residues, as well as the α-helices and β-strands of NAGK (purple), involved in the ligand coordination, are indicated.

The PII_ΔT-loop_ trimers that are bound to the NAGK hexamer, in turn, harbor three Mg^2+^-ATP molecules at the cleft between its trimer subunits (Figure 5C,E and Figure S3D). Mg^2+^-ATP is coordinated by surrounding T29, R94 and R96 from one PII subunit and R38 and K83 from the other PII subunit (Figure 5E). The overall structure of PII_ΔT-loop_ revealed a typical trimer with a ferredoxin-like fold similar to other known PII structures^1–3^ (Figure 5C and Figure S4C), however, lacking the extended T-loop (Figure 5C). In contrast to the previously published structures of PII_WT_ bound to NAGK^12,14,17^, the shortened T-loop of PII_ΔT-loop_ variant does not exhibit a characteristic β-hairpin-like conformation, which is protruding deeply into the NAGK cavity (Figure 5B,C, Figure S4C, and Figure S5A). Fragmentary density corresponding to the shortened T-loop suggests increased flexibility of this part of PII_ΔT-loop_ and did not allow us to build the model of the shortened T-loop unambiguously (Figure 6B).

**Figure 6.**
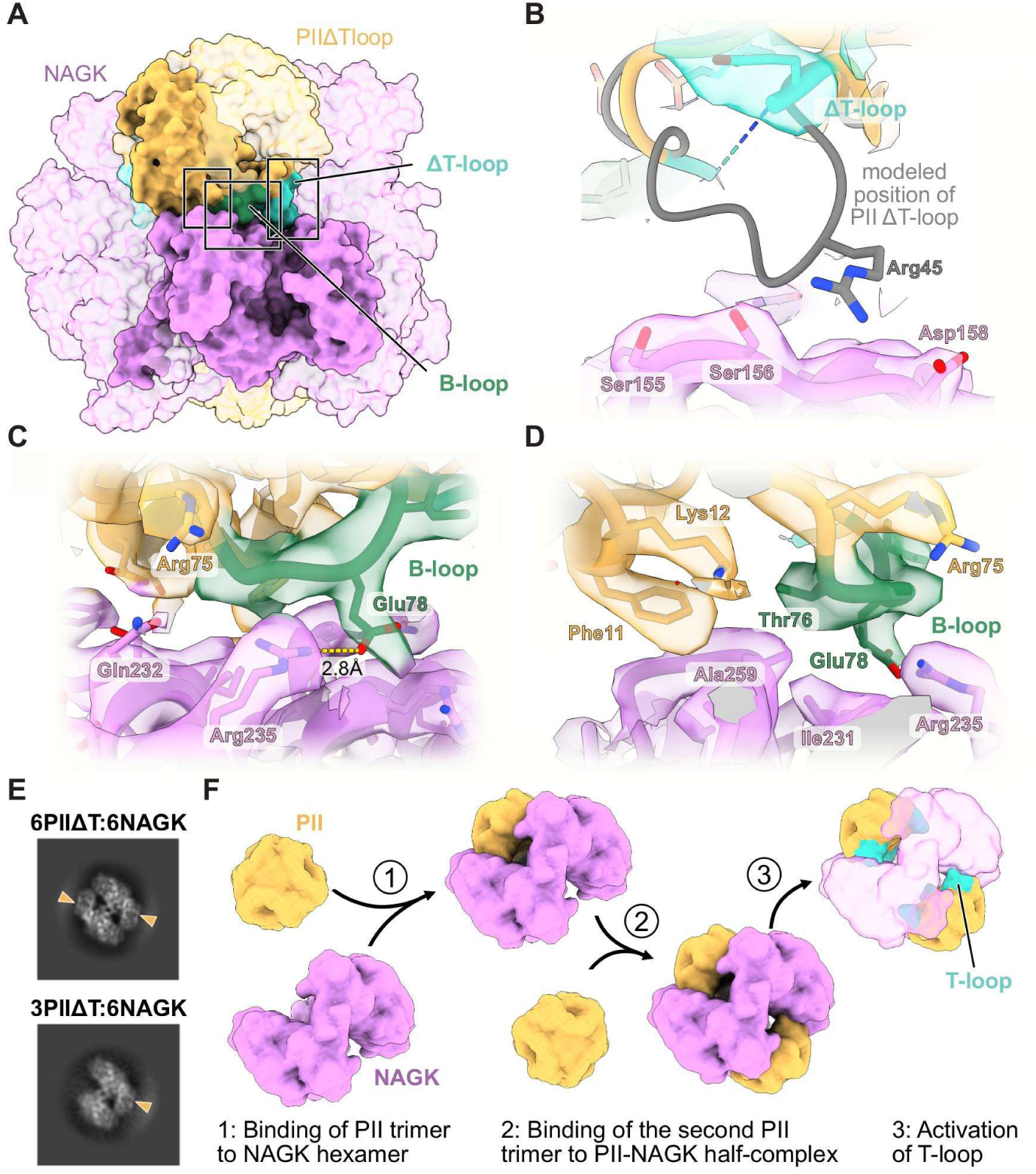
Interactions between PII_ΔT-loop_ and NAGK in the PII_ΔT-loop_-NAGK encounter complex. (A) Overview of the PII_ΔT-loop_-NAGK complex interface. The model is shown in surface representation (coloring is as in Figure 5). (B-D) Close-up views of the contacts between PII_ΔT-loop_ and NAGK (boxed in A). Truncated ΔT-loop in cyan (B), B-loop in green (C,D). The interacting amino acid residues are indicated. The associated cryo-EM densities are zoned around the models. The truncated T-loop is excluded from the final model due to the poor cryo-EM density quality in this region, and its hypothetical modeled position is shown in (B) in grey color. (E) Representative 2D classes showing a full PII_ΔT-loop_-NAGK complex (hexameric NAGK with two PII_ΔT-loop_ trimers attached, upper panel, yellow triangles) and a complex with only one PII_ΔT-loop_ trimer bound to NAGK hexamer (lower panel, yellow triangle). (F) Schematic model of PII-NAGK complex formation. Initially, NAGK hexamer binds to a PII trimer; after that, the second PII trimer binds to the assembly making the encounter PII-NAGK complex (as seen by our current model); finally, the insertion of the PII T-loop into the NAGK interdomain cleft initiates structural rearrangements within NAGK, including tightening the complex and bending of helix αN opening the Arg-binding site, which promotes NAGK activation and relieving from Arg-feedback inhibition.

The interface between PII_ΔT-loop_ and NAGK is established mostly through the contacts of the structured B-loop and neighboring residues of PII_ΔT-loop_ with the C-terminal domain of NAGK (Figure 6A-D). These connections involve the salt bridge between E78 of PII_ΔT-loop_ B-loop and R235 of NAGK and the hydrophobic contacts including F11 and T76 of PII_ΔT-loop_ and I231 and A259 on the NAGK side (Figure 6C,D). On the other hand, the shortened and partially disordered T-loop of PII_ΔT-loop_ does not insert into the NAGK interdomain cleft in contrast to homologous PII_WT_-NAGK structures (Figure S5A). The T-loop residue R45, which is crucial for maintaining NAGK in an active form, is displaced to the outside of NAGK in comparison with the structure of PII_WT_-NAGK complex (PDB: 2V5H).^14^ The poor map quality in the region of the truncated T-loop, probably caused by an increased flexibility of this part of the protein, did not allow us to unambiguously model the R45 residue of PII. However, according to our model, R45 would be too far from the NAGK D158 residue to form a salt bridge (Figure 6B).

In our structure, the NAGK ring exhibits a wide-open conformation with an approximate diameter of 107 Å, which is similar to the Arg-bound NAGK from *Thermotoga maritima* (approx. 104.6 Å) and in contrast to the Arg-insensitive NAGK from *Pseudomonas aeruginosa* (100 Å) (Figure S4A).^22^ In addition, the NAGK ring in our PII_ΔT-loop_-NAGK complex is much wider (approx. 104 Å) than the NAGK hexamers in the structures of PII_WT_-NAGK complexes (approx. 97-100 Å) (Figure S4B).^14,17^ Moreover, the N-helices that are essential for NAGK intersubunit contacts and located close to the Arg-binding site are in a straight conformation, similar to Arg-bound *T. maritima* NAGK (Figure S5C), while the analogous N-helices in Arg-free NAGK or PII_WT_-NAGK complexes displayed kinked conformations (Figure S5C,D). In our complex, the straight conformation of helix αN keeps the Arg-binding site closed and accommodating the bound Arg molecule bound (Figure 5D,G, S5C,D), thereby inhibiting NAGK. This result explains the sensitivity of the PII_ΔT-loop_-NAGK complex to Arg-feedback inhibition (Figure 4B) and further supports that a tight complex as seen in PII_WT_-NAGK structures (Figure S5) is needed to alleviate Arg inhibition, in agreement with our enzymatic assays (Figure 4). In our cryo-EM dataset, we also detected a subset of particles that resulted in a 2D class representing the NAGK ring bound to only one trimer of PII_ΔT-loop_ (Figure 6E). This conformation may represent an intermediate state preceding the encounter PII-NAGK complex, suggesting a sequential stepwise association of PII to the NAGK hexamer (Figure 6F).

## DISCUSSION

In the present study, we conducted a series of experiments to investigate the influence of PII (ATP, ADP and 2-OG) and NAGK (NAG and Arg) effector molecules on the PII-NAGK complex formation. For the first time, we showed a promoting effect for NAG on the complex formation (Figure 2) similar to the known ATP influence^15^ (Figure 1), while Arg showed antagonistic effect (Figure 2 and Figure 3) similar to ADP and 2-OG effectors of PII protein.^5,21^

Further, it was speculated that PII-NAGK complex formation follows a two-step mechanism, in which the B-loop of PII is initiating the interaction with NAGK in an encounter complex, followed by the bending of PII T-loop resulting in a conformation allowing its deep insertion into NAGK clefts.^16^ The deep insertion of PII T-loop into NAGK causes the complex to adopt a tight conformation, alleviating its Arg-induced inhibition and increasing NAGK activity.^14,17^ However, these assumptions have not been experimentally tested. Here, we examined this hypothesis by creating a PII variant, which lacks the T-loop and thereby blocks the complex in its first encountering state. We revealed that PII_ΔT-loop_ can still bind and activate NAGK, albeit with lower efficiency compared to PII_WT_. This demonstrates that PII can function partially and transmit signals to NAGK without the distal part of the T-loop. In line with this, we observed a weak density corresponding to the NAGK substrate NAG, indicating its partial occupancy in the complex. We also modeled ATP and not ADP in the corresponding density in the NAGK binding pocket, suggesting that the enzyme is trapped in a substrate-bound state, not in the catalytic active state. The PII_ΔT-loop_-NAGK complex was indeed blocked in the first association stage, where the interaction was mainly established through the PII B-loop residues (Figure 6C,D), particularly PII E85, which approaches NAGK R233, independently of the T-loop.^14^ In contrast, our data show that R45 of PII is not part of the PII-NAGK interface within the encounter complex (Figure 6B). R45 is known to act as an anchor residue via linking the T-loop to NAGK through an ion pair network and thereby increase the NAG substrate affinity to NAGK.^14,17^ However, in our cryo-EM map, the R45 density was too weak to allow its unambiguous modeling, suggesting this part of PII being flexible. Therefore, we propose that R45 does not coordinate the binding to NAGK, and may rather be involved in the deep T-loop insertion into NAGK after establishing the primary encounter complex. Consistent with that, the truncation of the nearby T-loop residues 47-53 caused a significant increase in the PII_ΔT-loop_-NAGK’s K_m_ value with a reduction in the NAG substrate affinity compared to the PII_WT_-NAGK complex (Figure 4). Our structural analysis also showed in general less contacts between PII_ΔT-loop_ and NAGK in the complex, in contrast to the wild-type complex and in agreement with the weaker binding kinetics (Figures 1-3).

In contrast, the interaction of PII_ΔT-loop_ with NAGK is not sufficient to narrow the NAGK ring to render it Arg insensitive (Figure S5) and maintains it in a wide conformation, which is similar to the previously published inhibited NAGK structure (PDB: 2BTY).^22^ Notably, the wide conformation and the associated straight conformation of the N-helix facilitates an increased affinity of NAGK to Arg, thereby enhancing the Arg-feedback inhibition (Figure S5C).^14,22^ Concurrently, in the NAGK wide-ring conformation, the nucleotide binding sites do not rearrange and remain far from NAG, therefore explaining the reduced NAGK activity in the PII_ΔT-loop_-NAGK complex compared to PII_WT_-NAGK (Figure 4). In contrast, when complexed with PII_WT_, the N-helix of NAGK bends and therefore causes the Arg-coordinating tyrosine residue (Tyr25) to move away and open the binding pocket allowing Arg to leave, thereby alleviating the Arg-caused feedback inhibition.

Since PII_ΔT-loop_ could not relieve NAGK from Arg-induced feedback inhibition, we suggest that multiple contacts between PII and NAGK and the insertion of the T-loop into the NAGK hexamer is required for tightening the NAGK hexameric ring and enhancing its activity. On the other hand, our findings confirm that the B-loop and the body of PII initiate the PII-NAGK complex assembly, leading to the formation of the encounter complex, while the PII T-loop is responsible for tight complex formation through anchoring deeply into the NAGK cavity. Interestingly, our cryo-EM analysis revealed a subset of particles representing NAGK complexed with only one trimer of PII_ΔT-loop_ (Figure 6E), which may represent the first intermediate state of PII-NAGK complex assembly. According to our model (Figure 6F), the association of PII with NAGK follows a sequential stepwise mechanism, in which initially one PII trimer binds to the NAGK hexamer, driven by the interactions provided by the PII B-loop, followed by the attachment of the second PII trimer and formation of the encounter complex. In the next step, the PII T-loops insert into the NAGK hexamer cavities, leading to a structural rearrangement in NAGK, including the tightening of the NAGK ring and a reorganization of the substrate binding sites. Such PII T-loop-induced conformational changes in NAGK alter its enzymatic kinetics. Firstly, the Arg-binding sites enlarge, decreasing the affinity of Arg for NAGK and thus relieving the Arg inhibitory effect. Secondly, the tight PII-NAGK complex boosts the affinity for the NAG substrate, thus enhancing the overall catalytic activity of NAGK.^11,14^ These results emphasize the importance of the B-loop in facilitating the formation of a stable encounter PII-NAGK complex (Figure 6E), independently of the T-loop.

Consistent with our findings regarding the B-loop’s importance, previous studies have surprisingly shown that a mutation around this region, as in the PII_(I86N)_ variant, causes constitutive binding to both NAGK and PipX^11,16^, the co-activator of the master nitrogen transcription factor NtcA.^23,24^ Unlike PII_WT_, the PII_(I86N)_ variant does not require positive stimulation by ATP or ADP to form stable complexes with NAGK or PipX, respectively. Furthermore, PII_(I86N)_ variant does not sense 2-OG^16^, therefore, the PII_(I86N)_-NAGK and PII_(I86N)_-PipX complexes do not readily dissociate in its presence.^11,16^ This causes the hyperactivation of NAGK^16,18^ with subsequent high Arg and cyanophycin (an Arg-based biopolymer) production^18^, and strong sequestration of PipX. These effects could explain the cyanophycin overproduction and the delayed nitrogen starvation response observed in the *Synechocystis* strain BW86 harboring the PII_(I86N)_ variant.^18^ Altogether, these findings underscore the critical role of the B-loop and the structural plasticity of the PII signaling protein.

## MATERIAL AND METHODS

### Bacterial strains and growth conditions

Recombinant PII_WT_ and PII_ΔT-loop_ from *Synechocystis* sp. PCC 6803, containing a C-terminally His-tag sequence^19^, were overexpressed and purified in *Escherichia coli* Lemo21(DE3) strain using the pET-15b vector system under the control of the T7 promoter^6,19^. Protein expression was induced with IPTG. Transformed bacteria were cultured in LB medium containing ampicillin (100 µg/ml) and chloramphenicol (34 µg/ml). Cultures were inoculated to an initial optical density at 600 nm (OD_600_) of 0.01 and grown at 37°C with shaking until OD_600_ of 0.6-0.8, then shifted to 20°C for overnight protein induction. Protein expression was induced by the addition of isopropyl-β-D-1-thiogalactopyranoside (IPTG) to a final concentration of 0.5 mM. Cells were harvested at an OD_600_ of approximately 3.0 by centrifugation at 3.500 rpm and 4°C for 15 min, and pellets were stored at-20°C. The gene encoding NAGK-Strep-tag from *Synechocystis* sp. PCC 6803 was cloned from previous plasmid^6,11^ into pET-15b vector, and overexpressed and purified from *E. coli* strain Lemo21(DE3) as described earlier for Strep-tagged proteins.^4,31^

### Cloning, expression, and purification of proteins

A gene coding for the truncated PII_ΔT-loop_ protein lacking amino acids residues (47-53) was synthesized (IDT, USA) and cloned into the pET-15b vector.^19^ To validate the construct sequence, the resulted plasmid was sequenced (Eurofins, Germany). Recombinant PII_WT_ and PII_ΔT-loop_ from *Synechocystis* sp. PCC 6803^19^, containing a C-terminally His-tag sequence, were overexpressed and purified in *E. coli* Lemo21 strain as described above.

For purification, cells were thawed at room temperature and resuspended in lysis buffer (10 mM imidazole, 150 mM NaCl, 100 mM Tris-HCl pH 8.6) supplemented with cOmplete Mini EDTA-free protease inhibitor cocktail (Merck, Germany). Cell debris was removed via centrifugation (Beckman Coulter Avanti J-26XP, USA) at 20,000 rpm and 4°C for 1 h. The supernatant containing the protein of interest was loaded onto a pre-equilibrated HisTrap™ column (1 mL, Merck, Germany). This was followed by extensive washing using wash buffers (150 mM NaCl, 100 mM Tris-HCl pH 8.0) containing stepwise increasing imidazole concentrations (10, 20, and 40 mM). Finally, protein was eluted using 500 mM imidazole. The eluate was collected and dialyzed against storage buffer (20 mM Tris-HCl pH 7.8, 0.5 mM EDTA, 5 mM MgCl2, 150 mM KCl, 100 mM NaCl, and 50% (v/v) glycerol) and stored at-20°C. Recombinant NAGK-Strep-tagged protein was overexpressed and purified as described previously.^4,31^

### Assay of NAG-kinase (NAGK) enzymatic activity

A coupled enzyme assay was used to determine NAGK activity in which the production of ADP was coupled to the oxidation of NADH by pyruvate kinase and lactate dehydrogenase as described previously.^15^ The reaction mixture consisted of 50 mM imidazole (pH 7.5), 50 mM KCl, 20 mM MgCl_2_, 0.4 mM NADH, 1 mM phosphoenolpyruvate (PEP), 10 mM ATP, 0.5 mM DTT, 5 U lactate dehydrogenase, 5 U pyruvate kinase, and 40 mM NAG, unless indicated otherwise. Where indicated, 0.5 µM PII_WT_ or PII_ΔT-loop_ (calculated as monomer concentration) was added to the mixture. The reaction was initiated by the addition of 0.05 µM NAGK (monomer concentration). The reaction time course was recorded for 10 min at room temp (≈ 25 °C) using a SPECORD 205 photometer (Analytik Jena) at 340 nm. The phosphorylation of one molecule of NAG corresponds to the oxidation of one molecule of NADH, indicated by a linear decrease in absorbance at 340 nm. One unit (U) of NAGK is defined as the amount of enzyme that catalyzes the conversion of 1 µmol of NAG per minute, calculated using the molar extinction coefficient of NADH (ε340= 6178 L mol^−1^ cm^−1^). The enzymatic parameters K_m_ and V_max_ were calculated using GraphPad Prism 6.01 (GraphPad Software, USA).^16^ The assays were performed in at least in triplicates to calculate the kinetics.

### Biomolecular Binding Kinetics Assay

*In vitro* binding studies were performed using Bio-layer interferometry (BLI) on an Octet K2 system (FortéBio), as described previously.^20^ Initially, His_8_-tagged PII_WT_ or PII_ΔT-loop_ variants were immobilized on Ni-NTA biosensors (FortéBio), followed by a 60 s baseline measurement. To analyze the binding of Strep-tagged NAGK, the loaded biosensors were immersed in NAGK solution (ranging from 31 to 1000 nM) for 180 s (association phase). To examine the influence of effector molecules (1 mM ATP, 1 mM ADP, 1 mM 2-OG, 50 mM NAG, or 100 µM Arg) on the PII-NAGK complex assembly and disassembly ^9,11^, these metabolites were added separately or in combination to the binding buffer as indicated. HEPES buffer (50 mM HEPES-KOH pH 8.0, 5 mM MgCl_2_, 150 mM KCl) was used for all experiments. Finally, dissociation was monitored for 300 s by transferring the biosensors into buffer alone at room temp (≈ 25 °C). For each experiment, a parallel reference sensor (negative control) without the interaction partner was included to correct for non-specific binding. Biosensors were regenerated after each run using 10 mM glycine (pH 1.7) and recharged with 10 mM NiCl_2_. Binding kinetics (V_max_) and the equilibrium dissociation constant (K_d_) were calculated using GraphPad Prism by fitting the obtained curves (GraphPad Software, USA).^20^ The binding assays were performed at least 2-3 times to calculate the binding kinetics.

### Size exclusion chromatography coupled to Multiangle Light Scattering

To determine whether the PII_ΔT-loop_ variant physically interacts with NAGK to form a stable complex similar to PII_WT_, Size Exclusion Chromatography coupled to Multi-Angle Light Scattering (SEC-MALS) was performed. Experiments were conducted at room temperature using an Äkta chromatography system connected to a Superdex 200 Increase 10/300 GL column (Cytiva). The system was coupled to a miniDAWN TREOS detector and an Optilab T-rEX refractometer (Wyatt Technology Corp.), as described previously.^9,32,33^ PII_ΔT-loop_ or PII_WT_ were analyzed separately or in combination with NAGK; NAGK alone served as a control. The column was equilibrated with running buffer (150 mM Tris-HCl pH 7.5, 150 mM NaCl, filtered and degassed), and runs were performed at a flow rate of 0.4 mL/min. For complex formation, PII variants and NAGK were mixed at a molar ratio of 3:1 (PII trimer: NAGK hexamer) and incubated for 10 min at room temperature prior to injection. Elution profiles were monitored by UV absorbance, and molecular mass was calculated from light scattering data using ASTRA software (Wyatt Technology).^11^ Subsequently, elution fractions were collected and analyzed via SDS-PAGE.^9^

### Cryo-EM sample preparation and data acquisition

Briefly, His-tagged PII_ΔT-loop_ and NAGK samples that were stored in the presence of 50% (v/v) glycerol as described above, were first cleaned using PD Minitrap™ G-25 (Cytiva) desalting columns at 4°C using buffer containing 50 mM Tris, 150 mM NaCl, 50 mM KCl, 5 mM MgCl_2_, pH8. Afterwards, samples were concentrated and further cleaned by SEC in the same buffer using a Superose 6 Increase 5/150 GL column (Cytiva) on an ÄKTA go purification system (Cytiva). After the SEC, the samples were mixed, resulting in final concentrations of 0.75 mg/ml for NAGK and 2.85 mg/ml for PII_ΔT-loop_. For cryo-EM, 2 mM ATP, 1 mM Arg and 10 mM NAG were added to the sample immediately before vitrification. 3 μl of the sample mixture were applied to glow-discharged C-Flat grids (R1.2/1.3 3Cu-50) (EMS) and directly plunge-frozen in liquid ethane using a Vitrobot Mark IV (Thermo Fisher Scientific) with the environmental chamber set at 100% humidity and 4°C. Movies (9149) were automatically collected using EPU (Thermo Fisher Scientific) on a Glacios cryogenic transmission electron microscope (Thermo Fisher Scientific) operating at 200 kV equipped with a Selectris energy filter and a Falcon 4 detector (both Thermo Fisher Scientific). Data collection was performed in Electron Event Representation (EER) mode at a nominal magnification of 130,000 (0.924 Å per pixel) in the defocus range of-0.8 to-1.8 μm, with an exposure time of 7.51 s, resulting in a total electron dose of approximately 52 e^-^Å*^−^*^2^.

### Cryo-EM image processing

The collected cryo-EM data was preprocessed in cryoSPARC Live, followed by additional processing in cryoSPARC v3 and v4^34^ (see Figures S2 and S3 for details). During preprocessing in cryoSPARC Live, motion correction, contrast transfer function (CTF) estimation, micrograph curation, and particle selection using a template picker (using templates generated from the structure of PII complexed with NAGK from *Synechococcus elongatus* PCC 7942, PDB: 2V5H)^14^, followed by 2D classification, were performed. Further steps were carried out in cryoSPARC. Briefly, the processing workflow involved selection of best micrographs (2392 micrographs, CTF-fit cutoff of 7 Å) followed by another round of particle template picking and two rounds of Topaz picking to select rare particle views. All picked particles were subjected to rounds of 2D classification, removal of duplicate particles and other rounds of 2D classification. Afterwards particles from best 2D classes were used for ab-initio reconstruction followed by heterogeneous refinement using 2 classes without applying a symmetry (Figure S3). The best class (92431 particles) was subjected to NU-refinement without applying a symmetry.^35^ Afterwards, particles were extracted with the full box size (0.924 Å per pixel) and subjected to another round of NU-refinement with D3 symmetry and defocus refinement. This resulted in a PII_ΔT-loop_-NAGK map at a resolution of 3.24 Å (Gold Standard Fourier Shell Correlation [GSFSC] value of 0.143). Unsupervised B-factor sharpening was applied and local resolution estimation was performed within cryoSPARC (Figure S2). Dataset statistics are provided in Table (S1).

### Model building and refinement

AlphaFold structure prediction of NAGK (Uniprot: P73326) and crystal structure of *PII (PDB: 1UL3) from Synechocystis*^26^ were manually fitted into the final cryo-EM map. The fragment of the T loop (residues 47-53) was removed from the structure of PII using Coot and the PDB-Tools Web interface.^36^ The models of NAGK and PIIΔT-loop underwent manual adjustments and refinements in Coot^27^, models of the ligands (Mg^2+^, ATP, Arg, NAG) were imported in Coot using the respective 3-letter code. Iterative rounds of real-space refinement in PHENIX^29^, accompanied by manual adjustments in Coot, were performed. Model validation was carried out using MolProbity^28^ in PHENIX (see Table S1 for model refinement and validation statistics). Visualization and figure preparation were performed in UCSF ChimeraX^30^ and Affinity Designer 2.

### Quantification and statistical analysis

The resolution of the cryo-EM map was determined using the gold-standard Fourier Shell Correlation 0.143 criterion, and local resolution estimates were generated in cryoSPARC. The quality of the atomic model was analyzed and validated with Coot, PHENIX and MolProbity. No methods were used to estimate sample sizes, and no blinding was done. Biochemical (NAGK enzymatic assays) and biophysical (BLI interaction experiments) data are presented as mean ± SD from at least 2-3 independent experiments. Statistical analyses and kinetic parameters for protein binding assays (V_max_ and K_d_) and NAGK activity assays (V_max_ and K_m_) were calculated using GraphPad Prism software. The statistical analyses were done as described in the figure legend.

## Data and code availability

Atomic coordinates of the PII_ΔT-loop_-NAGK complex have been deposited in the PDB under accession code 9TIQ. The corresponding cryo-EM map PII_ΔT-loop_-NAGK complex, has been deposited in the Electron Microscopy Data Bank (EMDB) under accession code EMD-55965.

## ACKNOWLEDGMENTS

The KAS laboratory is funded by the German Research Foundation - DFG as part of the SFB1381 (project number: 403222702; subproject B12) and Emmy Noether program (SE 3449/3-1). We also acknowledge the infrastructural support by CMFI (EXC 2124-390838134) and SFB1535 MibiNet (project number: 458090666). We thank Jörg Scholl (Tübingen University) for excellent assistance to create the plasmids used in this study. DS acknowledges the support by the DFG, including the Emmy Noether grant SH 1664/2-1 (project number: 537976353) and SFB1557. We thank Karl Forchhammer and Arne Moeller for continued support and constructive discussions, particular for access to the cryo-EM platform (DFG grant INST190/196-1 FUGG).

## AUTHOR CONTRIBUTIONS

KAS conceived, initiated, and supervised the research. KAS and AE designed the study. AE and DS performed research and prepared the figures. DS and KAS analyzed and interpreted the data. DS and KAS wrote the manuscript with inputs from AE. All authors approved the final version of the manuscript.

## DECLARATION OF INTERESTS

The authors declare no competing interests.

## Supplemental information

**Supplemental Figures**

**Figure S1.**
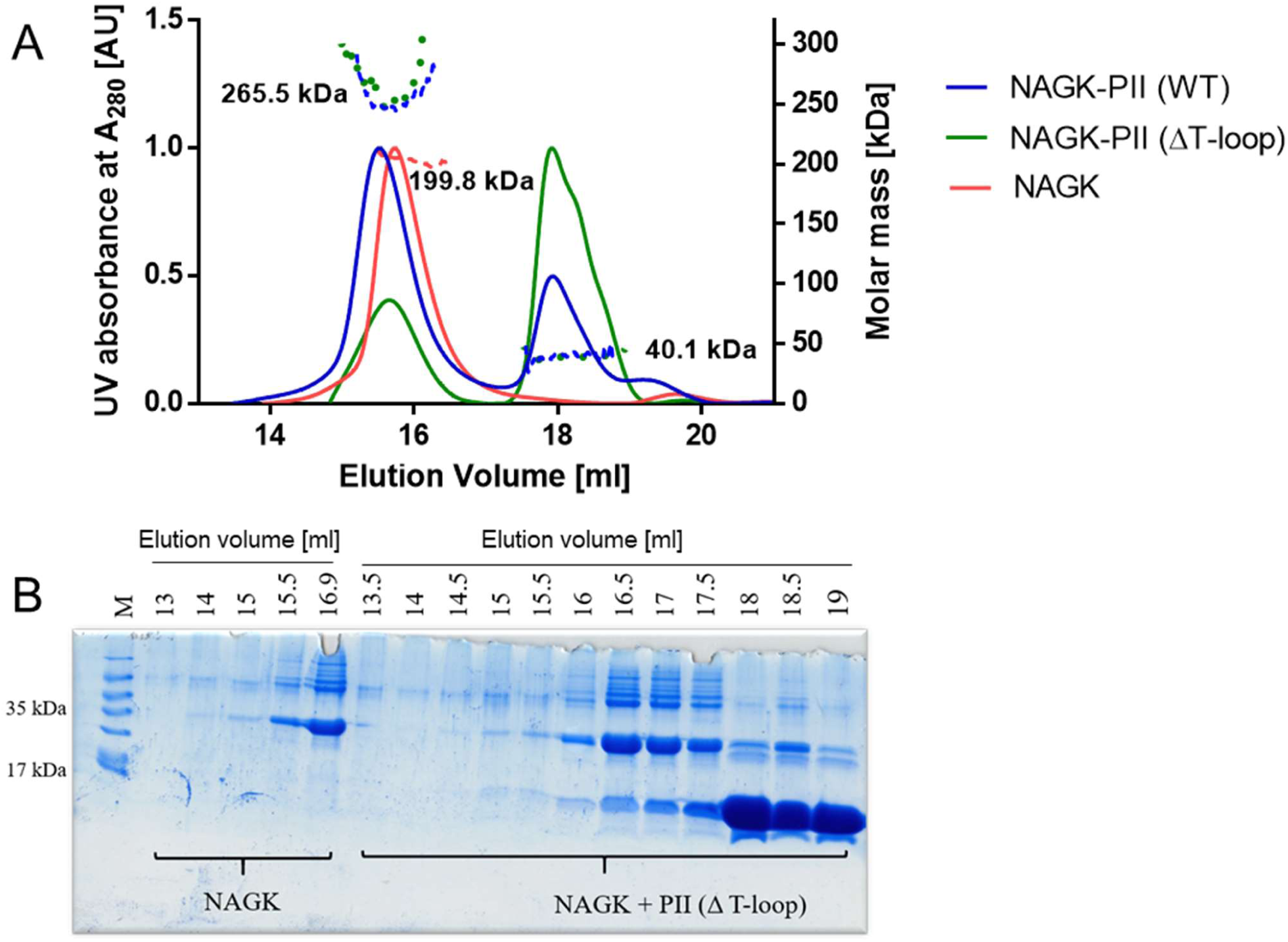
PII-NAGK complex formation with different PII variants using SEC-MALS. (A) SEC-MALS analysis indicates weaker complex formation between PII_ΔT-loop_ and NAGK compared to PII_WT_ with NAGK, despite similar molecular masses (blue and green dotted lines). (B) Eluted protein fractions in (A) were collected and subjected to SDS-PAGE followed by Coomassie blue staining. The SDS-PAGE analysis confirmed the presence of both PII_ΔT-loop_ and NAGK proteins in the PII_ΔT-loop_-NAGK complex peak. Top row: M, marker; 13-19, elution volume (ml) of fractions collected from (A).

**Figure S2.**
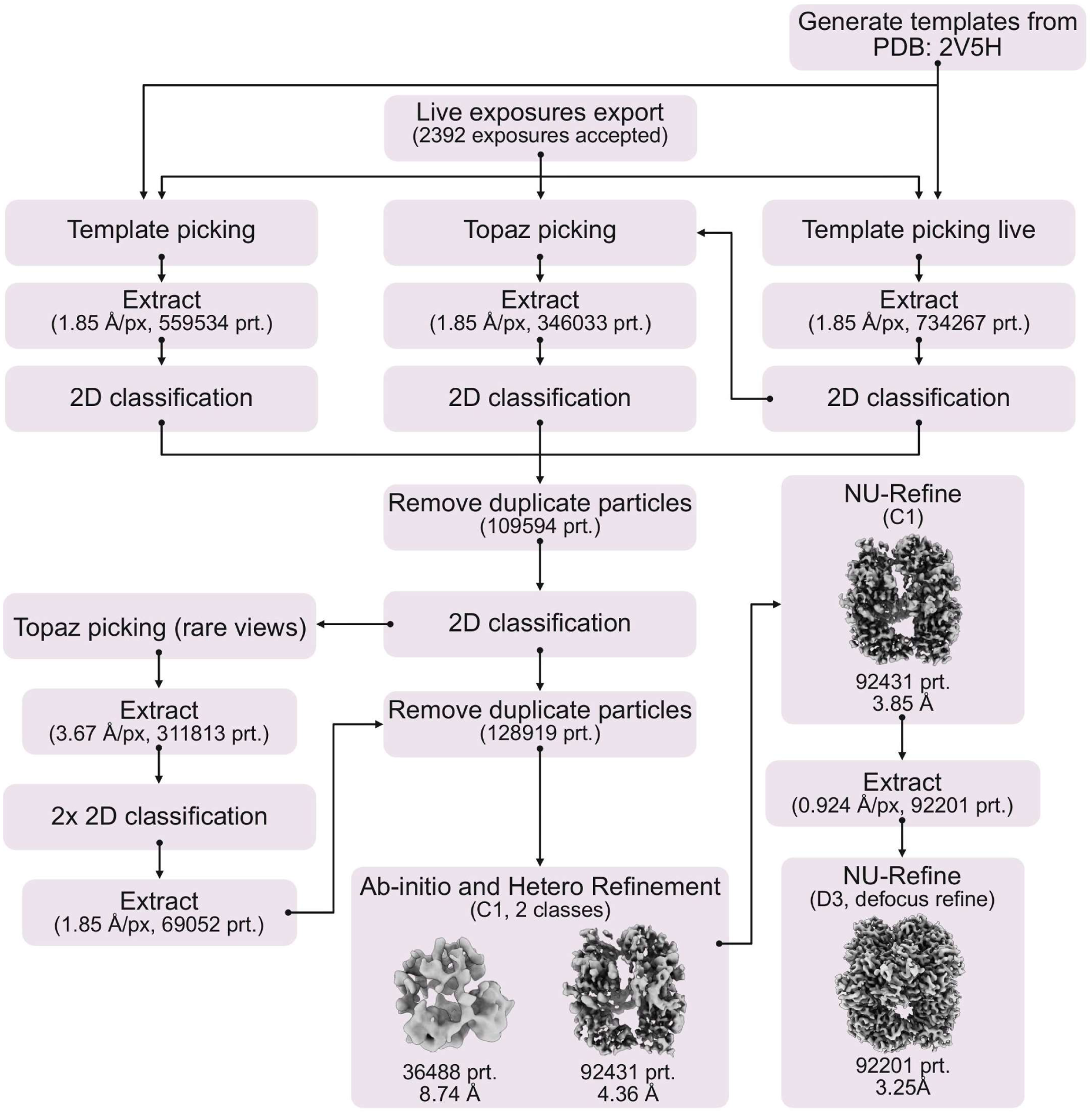
Cryo-EM image processing workflow. The key steps of data processing including the job types performed in cryoSPARC are shown. For extract jobs, pixel size and number of used particles are shown. For refinement jobs, applied symmetry, number of classes, number of particles, and achieved resolution are indicated.

**Figure S3.**
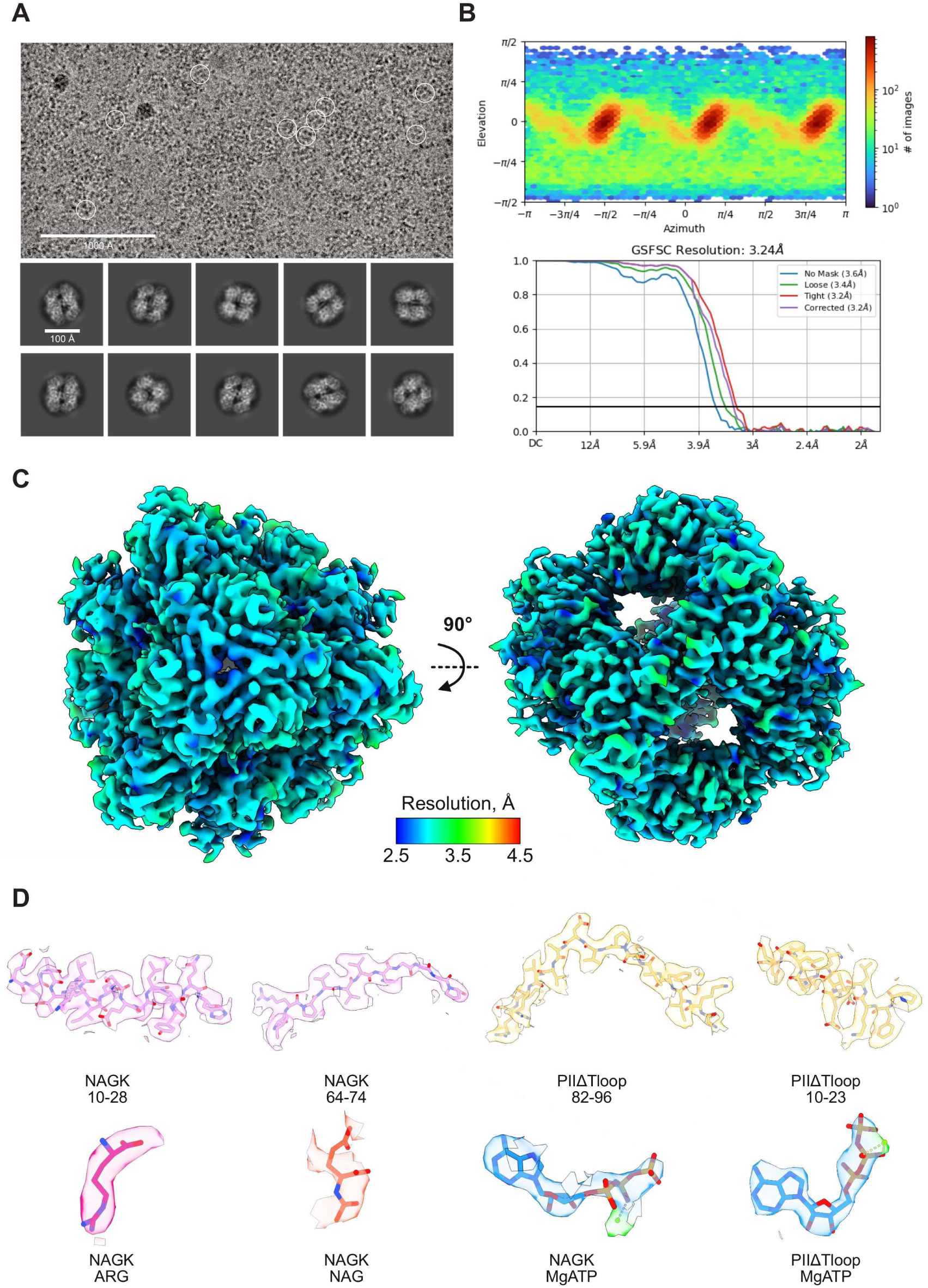
Cryo-EM data quality analysis. (A) Representative micrograph and 2D class averages depicting different views of the PII_ΔT-loop_-NAGK complex. Scale bars are indicated. (B) Angular distribution plot (top) and Gold Standard Fourier Shell Correlation (GSFSC) curve (bottom). (C) Local resolution estimation map of the PII_ΔT-loop_-NAGK complex. (D) Models of representative fragments of NAGK (purple) and PII_ΔT-loop_ (upper row), as well as ligands (bottom row) fitted in the corresponding cryo-EM densities (Arginine, Arg, pink; NAG, N-acetyl-L-glutamate, orange; ATP, dark blue, Mg^2+^, chartreuse).

**Figure S4.**
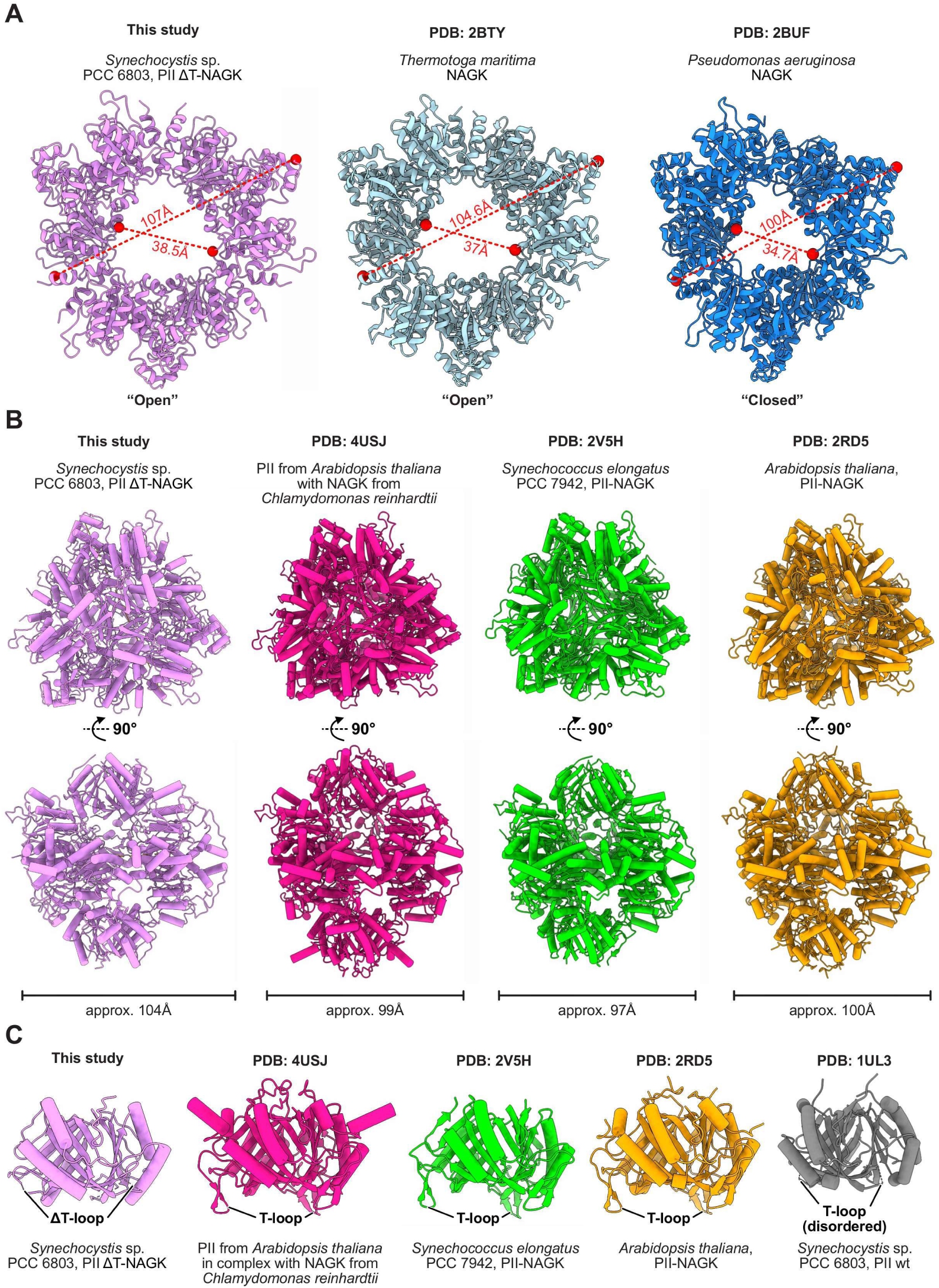
Structural comparison of the PII_ΔT-loop_-NAGK complex with homologous proteins. (A) NAGK hexamer from our PII_ΔT-loop_-NAGK structure viewed from the PII_ΔT-loop_-binding site (colored purple, PII_ΔT-loop_ model is hidden) compared with NAGK from *Thermotoga maritima* (PDB:2BTY, colored light blue) and from *Pseudomonas aeruginosa* (PDB:2BUF, colored dark blue). Outer and inner diameters of NAGK hexamer rings are indicated. (B) Structures of the PII_ΔT-loop_-NAGK complex from *Synechocystis* sp. PCC 6803 (this study, purple) and homologous PII-NAGK complexes from other organisms (PDB:4USJ, pink; PDB:2V5H, lime; PDB:2RD5, sand) viewed from the top and side. The approximate length of the complexes is shown at the bottom. (C) Trimeric structures of PII_ΔT-loop_ (this study, purple), PII_WT_ from complexes with NAGK (as in B; PDB:4USJ, pink; PDB:2V5H, lime; PDB:2RD5, sand) and free PII_WT_ (PDB:1UL3, grey) are shown. T-loop is indicated.

**Figure S5.**
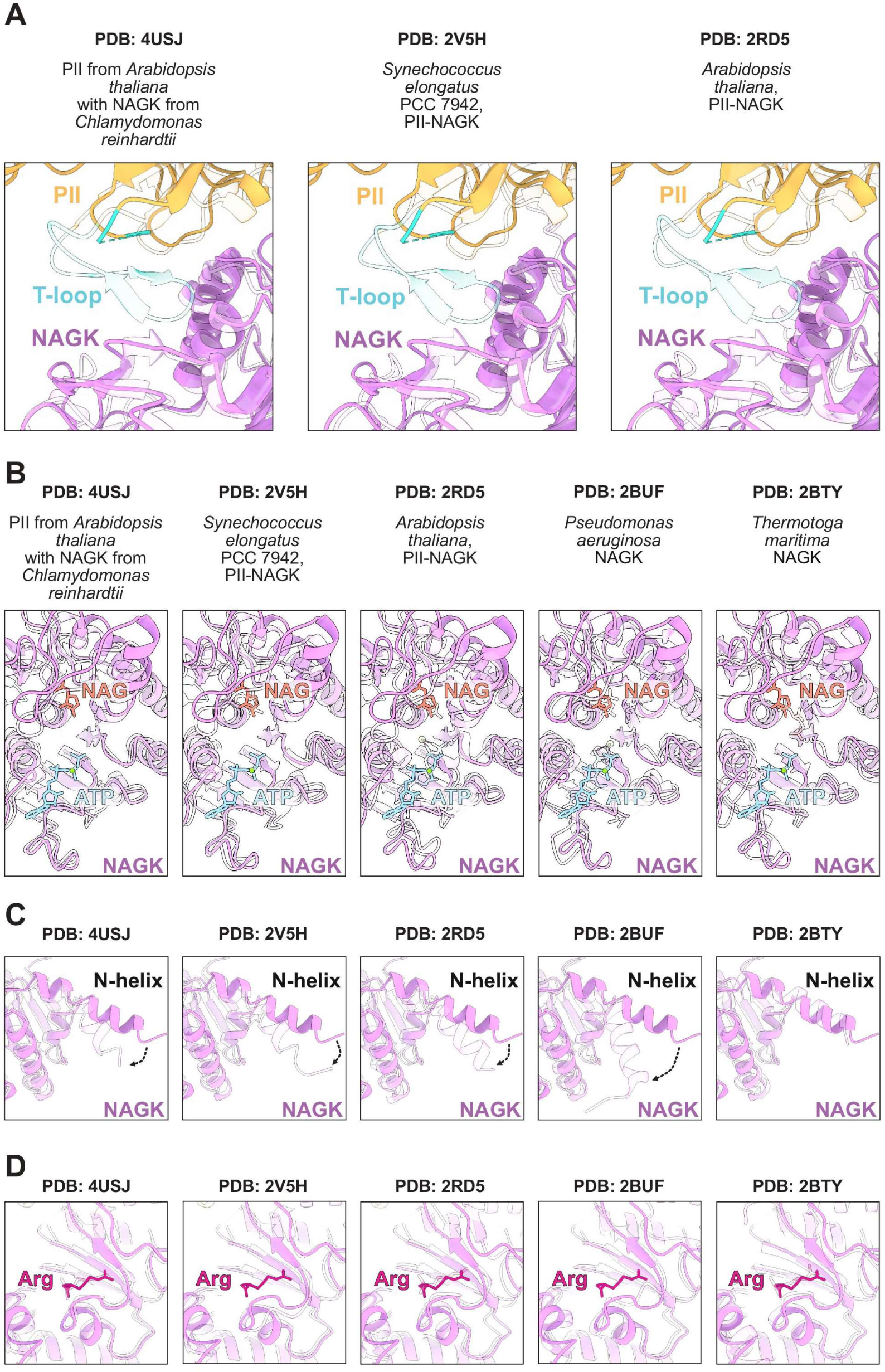
PII-induced conformational changes of NAGK. (A) The interface between PII (yellow) and NAGK (purple) demonstrating the characteristic T-loop (teal) conformation in the PII_ΔT-loop_-NAGK complex (this study, nontransparent) superimposed with the structures of homologous PII-NAGK complexes (semi-transparent). (B) Superposition of the NAGK (purple) active site from PII_ΔT-loop_-NAGK (this study, nontransparent) with the NAGK active sites from homologous structures (semi-transparent) of PII-NAGK complexes and free NAGK. Molecules of NAG (orange) and ATP (light blue) are indicated. (C) Superposition of the NAGK (purple) N-helix from PII_ΔT-loop_-NAGK (this study, nontransparent) with the analogous helix from homologous NAGK structures (semi-transparent). (D) Superposition of the NAGK (purple) Arg-binding site from PII_ΔT-loop_-NAGK (this study, nontransparent) with the analogous site from homologous NAGK structures (semi-transparent). The model of the Arg molecule is shown (pink). In (A-D), respective PDB codes of homologous published structures are indicated.

---

**Table S1.**
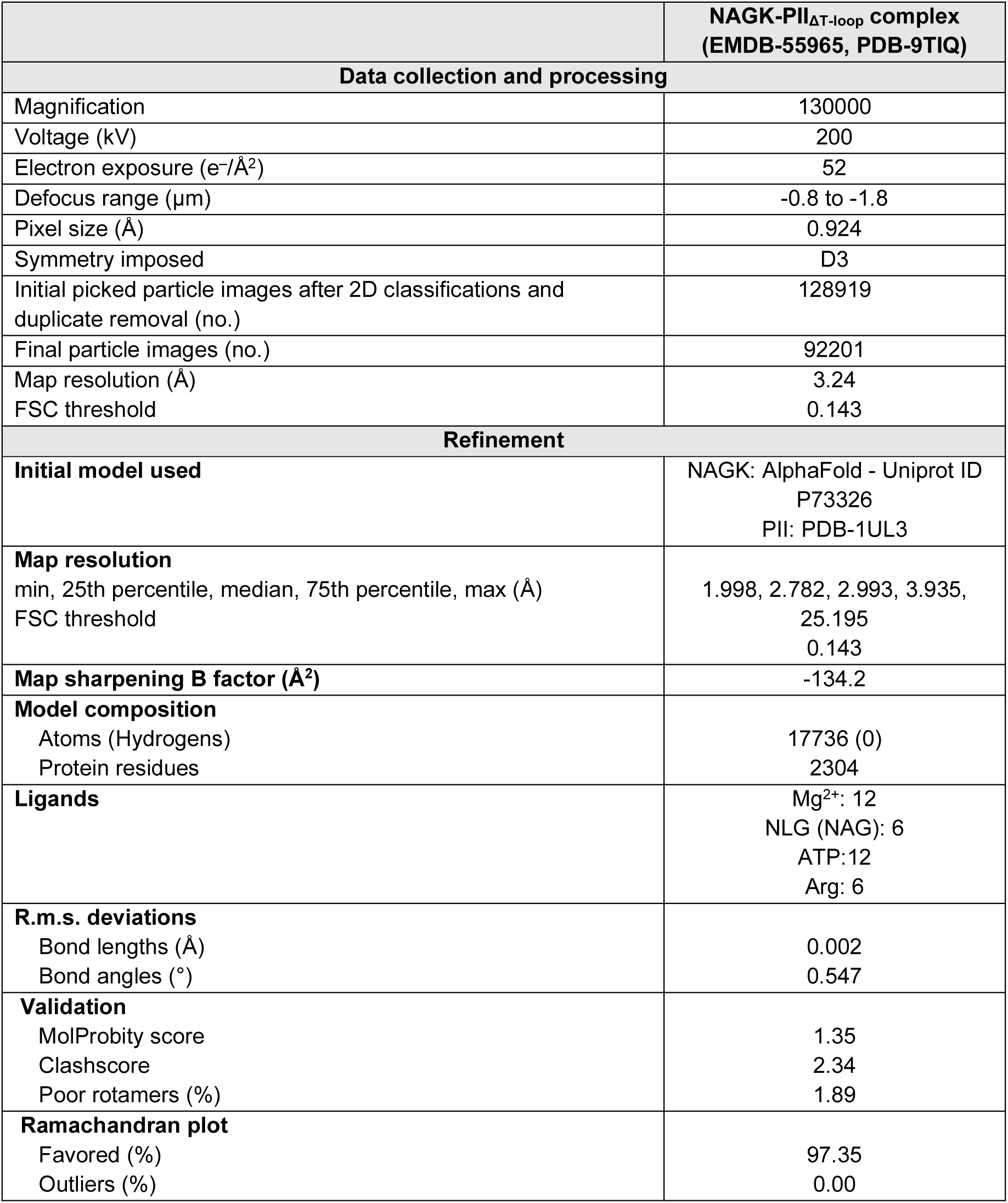
Cryo-EM data collection, refinement and validation statistics.

## Notes

### Competing Interest Statement

The authors have declared no competing interest.

